# Building containerized workflows using the BioDepot-workflow-builder (Bwb)

**DOI:** 10.1101/099010

**Authors:** Ling-Hong Hung, Jiaming Hu, Trevor Meiss, Alyssa Ingersoll, Wes Lloyd, Daniel Kristiyanto, Yuguang Xiong, Eric Sobie, Ka Yee Yeung

## Abstract

We present the BioDepot-workflow-builder (Bwb), a software tool that allows users to create and execute reproducible bioinformatics workflows using a drag-and-drop interface. Graphical widgets represent Docker containers executing a modular task. Widgets are then graphically linked to build bioinformatics workflows that can be reproducibly deployed across different local and cloud platforms. Each widget contains a form-based user interface to facilitate parameter entry and a console to display intermediate results. Bwb provides tools for rapid customization of widgets, containers and workflows. Saved workflows can be shared using Bwb’s native format or exported as shell scripts.

## Background

One of the key challenges for biomedical science is the rapidly increasing number and complexity of analytical methods. Reproducing the results of a bioinformatics workflow can be challenging given the number of components, each of which having its own set of parameters, dependencies, supporting files, and installation requirements. We present the BioDepot-workflow-builder (Bwb) as a solution to this problem. Bwb allows users to construct a graphical pipeline that connects different modules (widgets) together using a drag-and-drop interface. Instead of entering multiple command line flags, each software widget uses a form-based interface that allows users to enter and save parameters particular to that module. Users can edit a workflow by dragging a different widget onto the canvas and changing the linkages between the widgets. The resulting workflow can be executed in Bwb, saved in Bwb’s native format, or exported as a portable shell script that can be executed outside of Bwb.

Unlike other Docker based applications, Bwb supports graphical output and Graphical User interfaces (GUI’s), allowing for interactive tools such as Jupyter notebooks, spreadsheets and visualization tools to be included in the pipeline. Bwb’s workflows are portable with widgets that are self-contained and isolated from the operating system. Software updates of the host system or of individual widgets do not impact other widgets in the workflow. The modularity, portability and reproducibility come from Bwb’s use of individual Docker software containers for each of the software modules in the workflow. Most importantly, Bwb is open source and is distributed as a simple Docker container. Hence, Bwb can be easily deployed on any system (including Windows, Mac OS, Linux and any cloud provider) as long as Docker is installed. On cloud platforms, account management systems such as AWS Organizations can be used for sharing resources.

Software containers encapsulate each individual executable and its supporting libraries and software in isolated silos. This eliminates the possibility of conflicting dependencies and facilitates installation of software. Installed modules do not affect other modules or the host system. Version changes in any module are restricted to that container. The workflows constructed from containers are thus reproducible and portable. Docker is an application that manages these software containers on Linux, MacOS, Windows and cloud platforms [1]. Docker containers share the core hardware and operating system resources allowing them to be initialized in seconds. Docker containers can be built on demand using a small text file (Dockerfile) to define the components. Alternatively, Docker images can be downloaded from public repositories such as Docker Hub. If the required image is not available locally, Docker will even automatically download and install the image for the user.

Using Docker containers for individual modules in a workflow is an alternative to approaches that attempt to ensure that all components are compatible with a single computing environment. Software suites such as Bioconductor [2], BioPython [3], and BioPerl [4], provide a set of member components that are inter-compatible and ensure that the necessary dependencies for all components are installed. Ensuring that software is compatible within one of these frameworks is a non-trivial task. For example, Bioconductor requires that each component package compile and pass a suite of tests on Linux, MacOS and Windows platforms before each release. As a result, older software tools from research groups that lack the resources to publish, maintain and update their packages may eventually become excluded in future releases. Galaxy [5] takes the idea of a set of compatible modules a step further and provides a common web interface for users to create and execute workflows in a consistent hardware and software environment on a server or cluster. However, as is the case with Bioconductor, workflows may fail if all the components are not updated when Galaxy is updated.

As the numbers of software modules and dependencies continue to rise, the problem of some software components either failing or being deprecated with new releases increases. Workflows that depend on these legacy components must stay with older releases and older software or change to new unfamiliar software, which may break in future updates. As a result, it is very difficult for biomedical researchers to adopt state-of-the-art academic software. The problems associated with maintaining a set of compatible modules has led to increasing adoption of container-based tools such as Dockerstore [6], BioBoxes [7], BioShaDock [8], BioContainers [9], Bio-Docklets [10]. As additional examples, Bioconductor and Galaxy are also available in Docker containers [11, 12]. Packaging bioinformatics software in Docker containers has become increasingly common especially for cloud computing [13, 14] due to the ease of deployment on cloud instances.

Another important consideration for using Docker containers is the ease with which they can be stored without charge and made available for automatic download on Docker Hub [15]. In addition, many of these images are linked to Dockerfiles in GitHub source repositories [16] and rebuilt automatically as the code is updated. Public repositories now provide a readily accessible pool of containers for the construction of bioinformatics pipelines. BioContainers is one such repository currently harboring numerous Docker images [9] mostly based on automated bioconda build recipes. BioShadock [8] is another community driven registry of Docker-based bioinformatics tools [8] that also provides authentication, access control, container metadata, and search capabilities. Our group places all our containers in the BioDepot [17].

Other containerization efforts provide additional tools to facilitate the construction of workflows. For example, BioBoxes [7] uses a YAML Ain’t Markup Language (YAML) file to define the inputs and outputs of different containers [7]. NGSeasy, developed at the NIHR Biomedical Research Centre for Mental Health and Biomedical Research Unit for Dementia in London, UK [18], places all of the next generation sequencing (NGS) tools into one base image and encapsulates each NGSeasy pipeline component in separate containers. Dockstore, developed by the Cancer Genome Collaboratory [6], provides containers for their Toil scheduler [13] which supports the Common Workflow Language (CWL) for defining bioinformatics pipelines from the available component containers. As a demonstration, Toil was used to re-process, in parallel, 20,000 RNA-seq samples from RNA-seq data repositories such as The Cancer Genome Atlas (TCGA) and Therapeutically Applicable Research To Generate Effective Treatments (TARGET) on Amazon cloud instances comprising 32,000 computational cores in just four days [14]. As another example, Nextflow [19] provides scripting tools to design scalable and reproducible workflows using containers.

The most similar application to Bwb is Seven Bridges, a commercial service for conducting cloud-based bioinformatics analyses [20] which provides a GUI where users drag multiple “apps” onto a canvas and connect them together to define a workflow. The Cancer Genomics Cloud (CGC), powered by Seven Bridges, is one of the three platforms featured by the National Cancer Institute (NCI) Cloud Resources program [21]. However, Seven Bridges CGC consists of a web portal and is limited to the use of one cloud vendor (Amazon Web Services, AWS). On the other hand, Bwb is open source, available as a Docker container, and hence, can be deployed on any computer, any server and any cloud provider. Additional modularity is provided by parameters, customizable GUI entry forms, and Dockerfiles that are carried with the workflow themselves outside of the containers. Unlike Bwb, Seven Bridges CGC does not support graphical output for all applications. Finally, shared resources for a team must be set up via Seven Bridges. In contrast, Bwb is simply a Docker container and can leverage billing organizations provided by any cloud provider.

Galaxy [5] also provides similar functionality to Bwb and does have the ability to execute containerized workflows, for example, by using Bio-Docklets [10]. However, importing tools and containers from non-Galaxy sources is not trivial and requires modifying a set of configuration files and scripts whereas Bwb provides specific GUI tools for customizing existing workflows and can use containers from any of the Docker repositories. Export of workflows as bash scripts that can be customized and run without Bwb is also supported, whereas execution of Galaxy pipelines requires Galaxy.

## Results

### Overview

While the use of Docker alleviates many installation and reproducibility problems when deploying and executing bioinformatics workflows, several challenges remain. First, the command-line based interface used by Docker is not intuitive for biomedical researchers with limited programming experience. Secondly, software containers are designed for encapsulating software and software libraries and are not meant for storing parameters or data. While it is possible to hard code parameters and data inside software containers, this makes them unwieldy and difficult to customize. Third, Docker is designed for command-line pipelines and does not have graphical support out of the box. Useful interactive visualization and documentation tools such as Cytoscape [22] and Jupyter notebooks [23] can be difficult to incorporate into Docker pipelines. Finally, Docker containers can be quite difficult to customize, and versions of the containers can be difficult to track, especially with complex workflows. Bwb addresses each of these limitations as follows.

#### Bwb provides a friendly graphical interface

A graphical interface is provided through the use of our GUIdock-VNC [24] technology. Bwb is itself a container which is accessed through a browser. Instead of writing a script, the workflow is completely defined by the widgets and the graph that connects them. Construction of the workflow uses an intuitive drag-and-drop interface from the OrangeML library [25]. Links between widgets indicate the flow of data and order of execution. When a module has finished processing, it signals downstream modules to begin execution. Bwb translates this graph of widgets into a sequence of Docker commands which are then executed by the Docker engine. The complexity is all hidden from the user who merely clicks on a start button and observes the intermediate output on the widget consoles as the execution of the analytical pipeline progresses. See Figures 1 to 3 for examples of the Bwb graphical interface.

**Figure 1.**
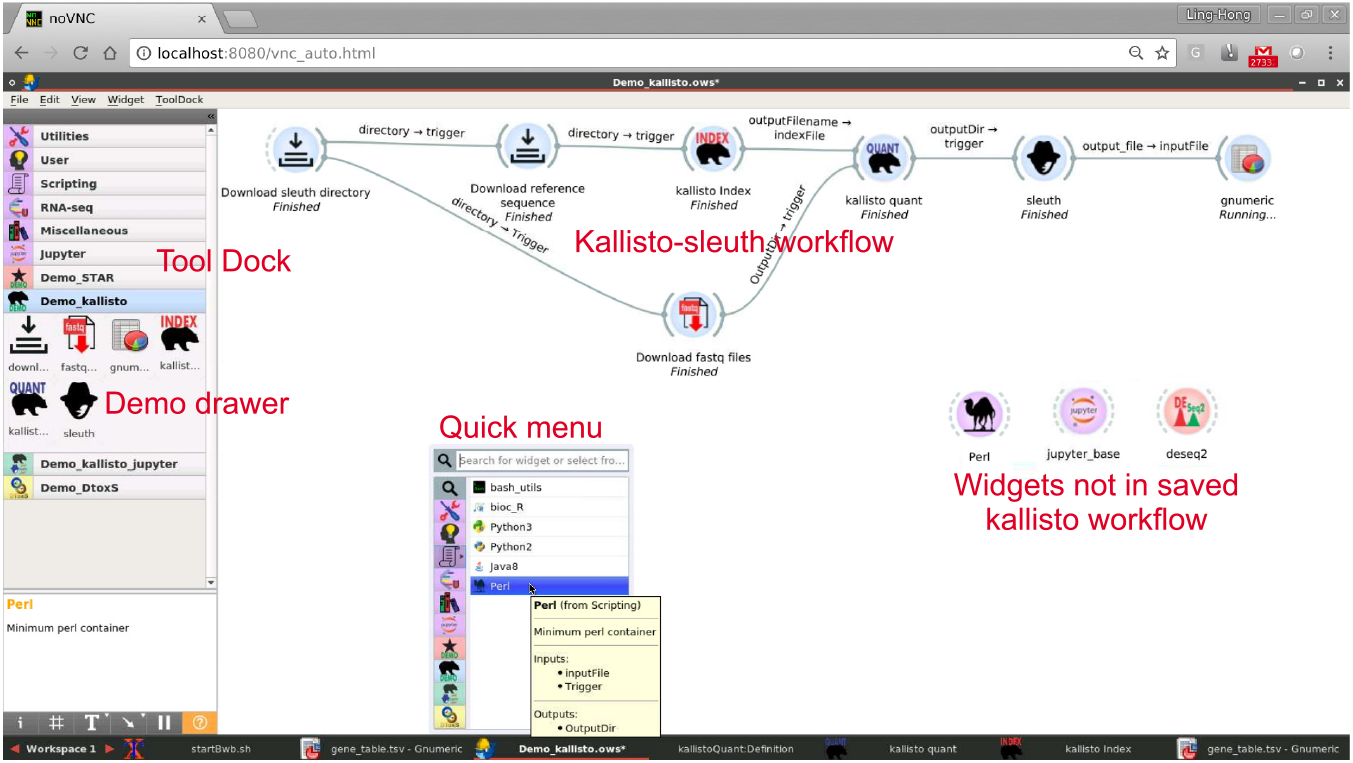
Screenshot of Bwb. The mini-windowing environment provided in the Bwb container is shown being accessed using the Chrome browser. When the Bwb application is launched, a window consisting of a canvas and a Tool Dock appears. The Tool Dock has multiple tabs which contain ‘‘drawers” of widgets from different categories and workflows. Widgets (the nodes) are dragged from the Tool Dock onto the canvas and linked together to form a workflow, in this case, the Kallisto-sleuth workflow. Widgets can also be added to the canvas by right-clicking on an empty area, which brings up a Quickmenu version of the Tool Dock. When a workflow is opened, its widgets are imported into separate drawer the tool dock. New widgets can be dragged onto the Tool Dock to be mixed and matched into the workflow. These widgets are denoted by different colors. In this case, the purple Perl and Jupyter widgets have been dragged from the standard purple Bwb drawers of the tool dock. The pink deseq2 widget has been dragged from the STAR drawer

#### Bwb widgets promote reproducibility by using containers and facilitating data entry and local file mapping

While the use of software containers virtually eliminates variation due to dependencies, the reproducibility of results depends on more than just the software. Hidden parameters and configuration files are also major factors contributing to the irreproducibility of results. A common solution is to include parameters and configuration files in the container itself. However, this strategy makes it difficult to re-use and customize the container. The best practice to maximize the portability of software containers is to have data files and parameters outside of the container as much as possible. Bwb provides form-based user interface for parameter values, which are stored outside the container, in a human readable XML file that is saved with the workflow. Data files are not included in the container but are obtained from the local host files. Since manually mapping local files and directories to the internal Docker file system can be an error-prone process, Bwb uses a simple default mapping system using the mounted volume that is defined when Bwb was launched. The user can be queried to mount additional volume mappings if desired, but the default mapping is sufficient for most use cases. This means that the user usually does not have to mount volumes to access local files. Bwb automatically handles the mapping of filenames transparently so that the user can choose files by simply clicking through a directory tree without worrying about which path will be used within each container

#### Bwb widgets are easily customizable

Bwb provides tools to allow users to easily create and customize new widgets. Right-clicking and choosing ‘edit widget’ brings up screens that allow the user to define which parameters are queried, which volumes and ports are to be mapped, which command should be run and the Docker container to be used. Parameters are entered using a form based interface and Bwb auto-generates the widget user interface (UI) based on the form entries. See Figure 2 for details of the widget panels. To facilitate the construction of Docker containers we also include a tool called BiocImageBuilder [26]. In addition, Docker files that specify container images can be optionally stored with the widget definition to further facilitate customization of the widget container. The widget definition is stored in 3 human readable JavaScript Object Notation (JSON) files.

**Figure 2.**
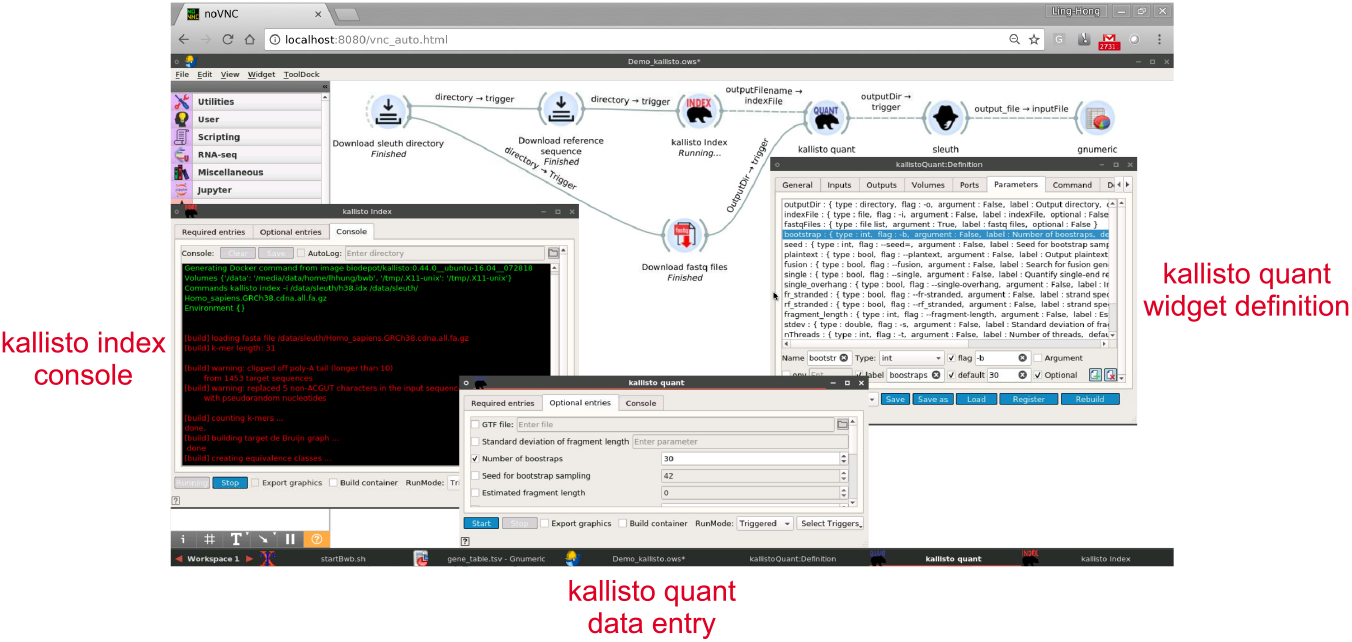
Widget panels. Each widget has two tabbed panels. Double-clicking on the kallisto-quant widget in the kallisto-sleuth pipeline brings up a data entry panel with a series of tabs. The numerous optional parameters that control the way that kallisto performs its quantifications are exposed by clicking on the ‘optional’ tab. Right-clicking the kallisto-quant widget and choosing edit brings up the widget definition panel. This reveals the settings that define the widget itself. The blue highlighted selection in the parameters tab of the kallisto:definition window shows the parameters defining the number of bootstraps option. The user can enter new values into the definition:window to change the default number of bootstraps for example. Finally, the black background window on the left is the console of the kallisto index. It displays the messages being printed by the widget as it processes the data. The data remains in the window for review until cleared.

#### Bwb workflows are encapsulated, portable, shareable and reproducible

In Bwb, workflows are defined by a graph of widgets and their parameters. Natively, Bwb stores workflows as a directory of widgets, and an XML file that describes the connections and the parameters. As long as the containers used are available on in a repository such as Docker Hub or defined in the widgets Dockerfiles, the workflow directory contains a complete description of the workflow logic, with the possible exception of parameter data files. However, even data files can be encapsulated using download widgets as shown in the case studies. Workflow directories contain human readable files and can be shared with other Bwb users. Finally, workflows can also be exported as a bash script consisting of the Docker commands that Bwb would execute on the host system. This script can be run without launching Bwb though there are some caveats as to GUI export and filenames as detailed in the Discussion section and in the user manual (Additional File 1).

#### Bwb supports export of graphics and GUI’s

Each widget contains a console tab that displays text output allowing the observation and logging of intermediate results. However, it is sometimes desirable to pop-up a window and to provide additional graphical feedback or to prompt user interaction. By checking a box, Bwb allows the widget to export graphics to the browser window using the methodology described in GUIDock-X11 [27]. This allows for commodity applications such as gnumeric (the gnu version of Excel) and Jupyter that pop up their own windows and have a their own GUI to be containerized and made part of the pipeline. Graphical applications such as Jupyter notebooks are especially useful for interactively monitoring the progress of execution or for customizing analyses and visualization of the final results. See Figure 3 for an illustration of graphical output in Bwb.

**Figure 3.**
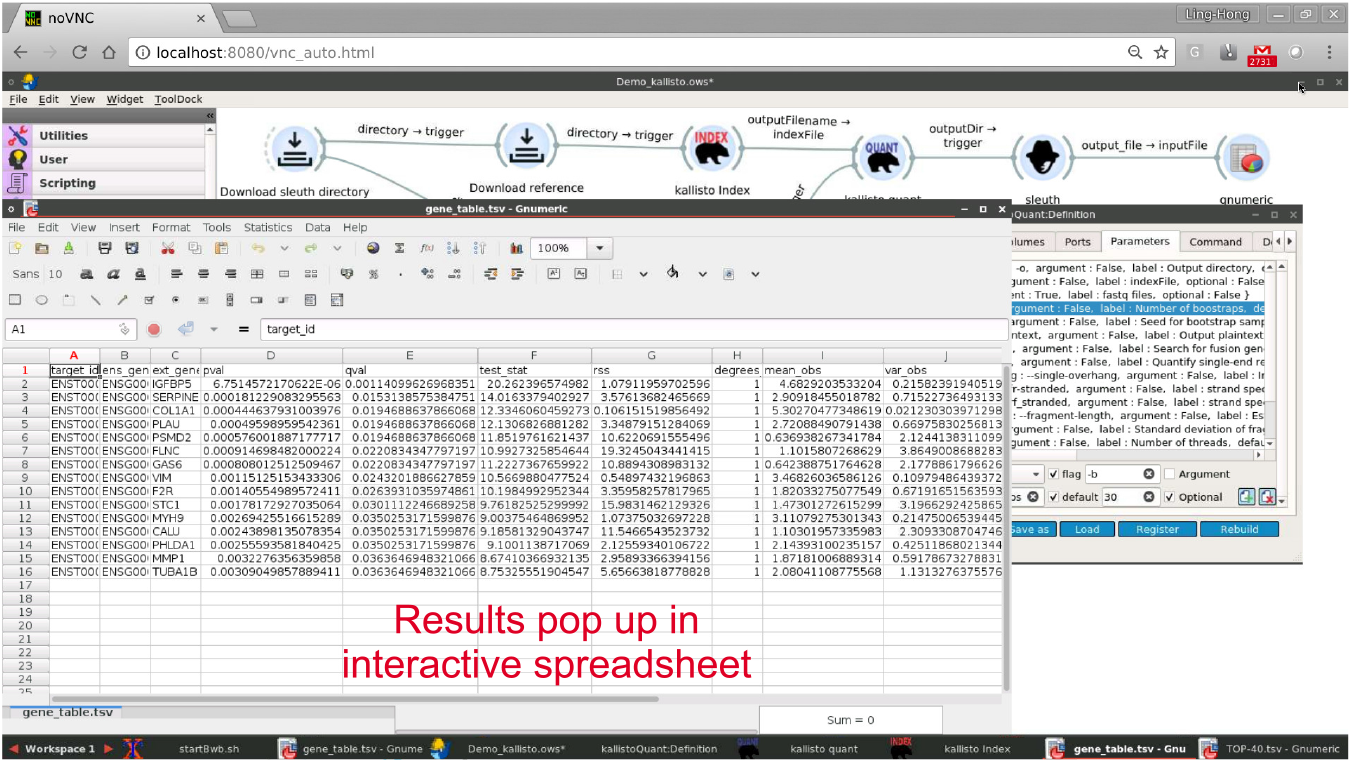
Graphical output of kallisto-sleuth workflow. Sleuth has its output linked to the trigger of the gnumeric spreadsheet. When sleuth is finished processing it sends the output to the trigger which prompts the gnumeric application to read the output CSV file and display the results. The window that is popped up is a normal gnumeric window and the user can interact with the process and visualize the data.

#### Bwb adheres to the FAIR principles

We designed Bwb with core values that are compliant with the FAIR Guiding Principles [28]. While the high-level FAIR Guiding Principles were primarily designed to support the reuse of scholarly data, these principles define characteristics that are also applicable to tools and infrastructures to facilitate scientific discovery and reuse. In particular, Bwb demonstrates *findability (“F”)* by exploiting ability of Docker to automatically download images from repositories. Bwb is able to use software components that are readily available in our BioDepot repository and other searchable container repositories. Bwb also provides tools to search and create containers from the CRAN (Comprehensive R Archive Network) [29] and Bioconductor [2] repositories. Bwb itself is easily findable through GitHub and Docker Hub which provides persistent record of current and past versions of Bwb source code and implementations. Bwb showcases *accessibility (“A”)* by providing a customizable drag-and-drop user interface such that scientists with all levels of technical expertise can access and interact with big biomedical data and software tools. Bwb uses Docker containers to encapsulate the computing environment for each task in bioinformatics workflows, thus ensuring interoperability (“I”) and re-usability (“R”). In addition, the entire workflow can be saved or exported, either as a template to be run with different data or with the original data for a complete and reproducible description of an analytical procedure, further encouraging re-usability.

### Case studies

We demonstrate the utility of Bwb using 4 case studies. The first case study is an example using Bwb to document and disseminate an existing workflow. The second and third examples illustrate the use of Bwb in well-established RNA-seq workflows using kallisto-sleuth [30, 31] and STAR [32]. In our final case study we demonstrate how to integrate Jupyter notebooks into Bwb. The combination of documentation, interactive dynamic code and graphics in Jupyter notebooks adds highly desirable functionalities to a workflow building tool. We demonstrate these functionalities by attaching a Jupyter notebook to the kallisto pipeline to run sleuth. Furthermore, we provide a tutorial and video showing how to add a Python script to this pipeline to trim the reads before processing (see video Additional file 3 in Supplemental materials) The saved workflows and widgets from these case studies are publicly available from GitHub and are included with the Bwb container. After downloading the Docker image of Bwb, users can simply load these case studies into the canvas and execute these workflows with one click.

#### Example 1: Reproducibly disseminating a Standard Operating Procedure using Bwb

The first case study is taken from the NIH-funded Library of Integrated Network-Based Cellular Signatures (LINCS) program which provides large-scale expression data from cell lines in response to genetic and drug perturbations [33]. The Drug Toxicity Signature Generation Center (DToxS), one of the LINCS data generation centers, studies the expression of human heart muscle cells under the influence of different drugs. As part of the effort to document all procedures and increase their reproducibility, laboratory and computational protocols for all experiments performed by DtoxS are given in detailed standard operating procedure (SOP’s) available on their website [34].

One of these SOP’s describes a RNA-seq data analysis workflow consisting of two major steps – alignment using the Burrows-Wheeler Aligner (BWA) [35] and quantification of differential gene expression using the R package edgeR [36]. We use this DToxS RNA-seq pipeline as a test case for the creation and dissemination of reproducible workflows using Bwb. This is an example of a SOP represented as a Bwb workflow. It is a completely reproducible, executable and sharable representation of the pipeline.

Figure 4 shows a demo of the DToxS RNA-seq workflow that can be run in a few minutes. This workflow consists of 5 widgets. The first widget (downloadURL) downloads the data and parameter files and sets up the directory structure. Note that the widget has been renamed to provide more clarity as to the role in the workflow. Renaming does not affect the program logic. The widget runs a custom bash script that calls the Linux curl utility to download gzip or bzip2 to decompress the files. The bash script also properly recognizes Google-drive URL’s. The docker image uses the `biodepot/bash-utils:alpine-3.7` container which is simply bash with wget, curl using the lightweight Alpine Linux operating system. This widget connects the output to the trigger of the next widget so that the second widget will start once the download is complete.

**Figure 4.**
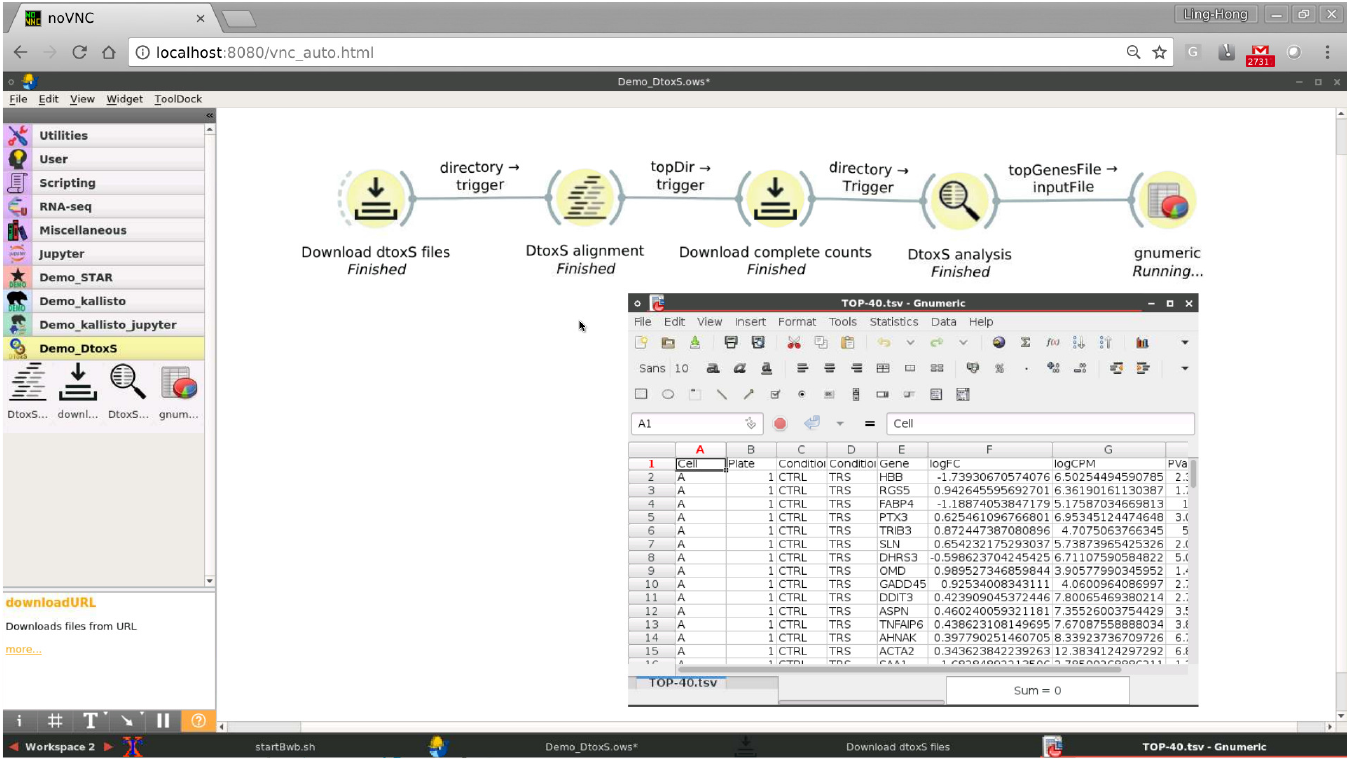
Screenshot of DToxS workflow demo. The DToxS RNA-seq workflow is implemented as a demo for Bwb. The connected workflow consists of 5 connected widgets and processes RNA-seq fastq files to obtain a list of differentially expressed genes. This list is displayed using the gnumeric spreadsheet

The second widget is the DtoxSAlignment widget. Again based on Alpine Linux, the container contains the BWA aligner and Python2.7 as these are necessary for the original alignment script. The output is connected to the trigger of the third widget to signal it to start once the alignment step is complete.

The third widget is another downloadURL widget. Normally we would connect the DToxSAlignment widget to the DToxSAnalysis widget. However, this pipeline on complete files takes over 12 hours and consumes 350 GB of space. For the demo, we use partial input files with fewer reads so that the alignment step can be completed in a few minutes. This will lead to an error in the subsequent analysis steps as there are too few counts. So for the demo, we download the complete results of the alignment using this widget. Again the output is connected to the trigger of the fourth widget. The fourth widget analyzes the gene counts obtained from the alignment step to determine which genes are differentially expressed. The script runs under R and uses Bioconductor. The script and these supporting packages are installed in a Ubuntu based image. In this case, when the widget is finished, it not only transfers a trigger signal but also the name of the output file in the format of CSV (comma-separated values) to the last widget so that it also knows which file to open.

The final widget is gnumeric, a fully functional open-source spreadsheet that duplicates many of the functions of Excel. It is inside a Ubuntu container and exports its own window and graphics. Note that the export graphics box in the widget options panel is checked to enable this functionality. Gnumeric displays the genes which are most likely to be differentially expressed (lowest p-values).

To start the demo, the user can double click on the first downloadURL widget and the start button. Clicking on the console tab reveals the output. As each widget finishes and triggers the next widget, we can follow the intermediate progress by double clicking on widgets and examining the console. After about 5 minutes, the final results should automatically pop up in the gnumeric spreadsheet which is shown below the workflow in Figure 4. To save the workflow as a bash script, the user checks off the ‘Test mode’ checkBox in the downloadURL widget before pressing start. Instead of executing the Docker commands generated by the workflow, Bwb will print the docker commands on the console of the widgets and optionally save the set of commands as a script.

#### Example 2: Kallisto-sleuth pipeline

In this second example, we illustrate the use of Bwb with a widely-used kallisto-sleuth RNA-seq data processing pipeline. The kallisto pipeline is shown in Figures 1–3. Kallisto takes the reads in fastq files and quantifies them using a rapid pseudoalignment technique [30]. Sleuth then processes the pseudocounts to find the genes that are differentially expressed [31]. This workflow is adapted from the introductory sleuth-walkthrough from the Pachter group [37].

The first widget again is a downloadURL widget that sets up the base directory and downloads two configuration files. One file defines which files are from the control and which files are from the treatment groups. The other file is used to map transcript names to gene names. The method of downloading configuration files rather than including them inside the containers makes the logic clearer and the workflow more easily customizable. This workflow bifurcates into two branches: one responsible for creating the indices and the other for downloading the fastq input files. The lower branch uses the widget created using the fastq-dump utility from the SRA toolkit to download the fastq files from the SRA (Sequence Read Archive) repository [38]. By default, the download is set to 10,000 spots, instead of the complete files, so that the demo can be completed in 20 minutes. The upper branch of the workflow consists of a download widget that fetches the human reference sequence and passes it to the kallisto index widget which produces the indices. The two branches merge at the kallisto quant widget which waits for both the indices to be made and the files to be downloaded before it can start. Sleuth then analyzes the counts created by kallisto quant and determines which genes are differentially expressed. The table of differentially expressed genes is passed to the gnumeric widget which pops up an interactive spreadsheet with the final results. The kallisto widgets are based on the executable compiled from the source in the GitHub repository. We could have used one widget for both the index and quant functions but the workflow logic is clearer with two different widgets and the kallisto quant widget required a wrapper script to handle multiple samples. Sleuth also uses a shell wrapper to pass parameters to an R script. The other widgets used containers that were straightforward installations of existing software, and using the widget builder to pass the flags and options to the user.

This workflow illustrates one of the key advantages of using Bwb – reproducible and automatic installation. Sleuth can be challenging to install requiring different (undocumented) supporting libraries and packages depending on the version of sleuth, the operating system and the version of R and whether R is being run with Jupyter. Using Bwb, the containers with all the necessary dependencies are automatically downloaded, and run identically regardless of the user’s host setup. Although the installation details are masked, they are accessible in the accompanying Dockerfile.

#### Example 3: STAR pipeline on a remote server

The STAR/DESeq [32, 39] pipeline in Figure 5 is another well-established workflow for identifying differentially expressed genes from RNA-seq experiments. The structure of the pipeline is similar to the kallisto pipeline and the construction of widgets similar, with a wrapper around STAR quant [32] to handle multiple samples. Again, the workflow is fairly self explanatory with the entire workflow being present. With STAR, there are literally pages of flags (shown in part in the scrollable panel in Figure 5) with many different options so the Bwb’s use of a form based interface to modify parameters is a very useful feature. STAR requires a large amount of RAM (a minimum of 32 GB is recommended) and the generation of indices and alignment steps are difficult to run on local hardware. This example is actually run on a remote firewalled server that is accessed securely through an ssh tunnel. The gnumeric spreadsheet pops up and works exactly the same as the other examples run locally on a laptop. However, even without a larger server, one can start the workflow at later stages as long as the necessary files are in place. The rest of the pipeline with parameters in place for documentation, is executable if the user wishes to use a larger local or cloud server at some point.

**Figure 5.**
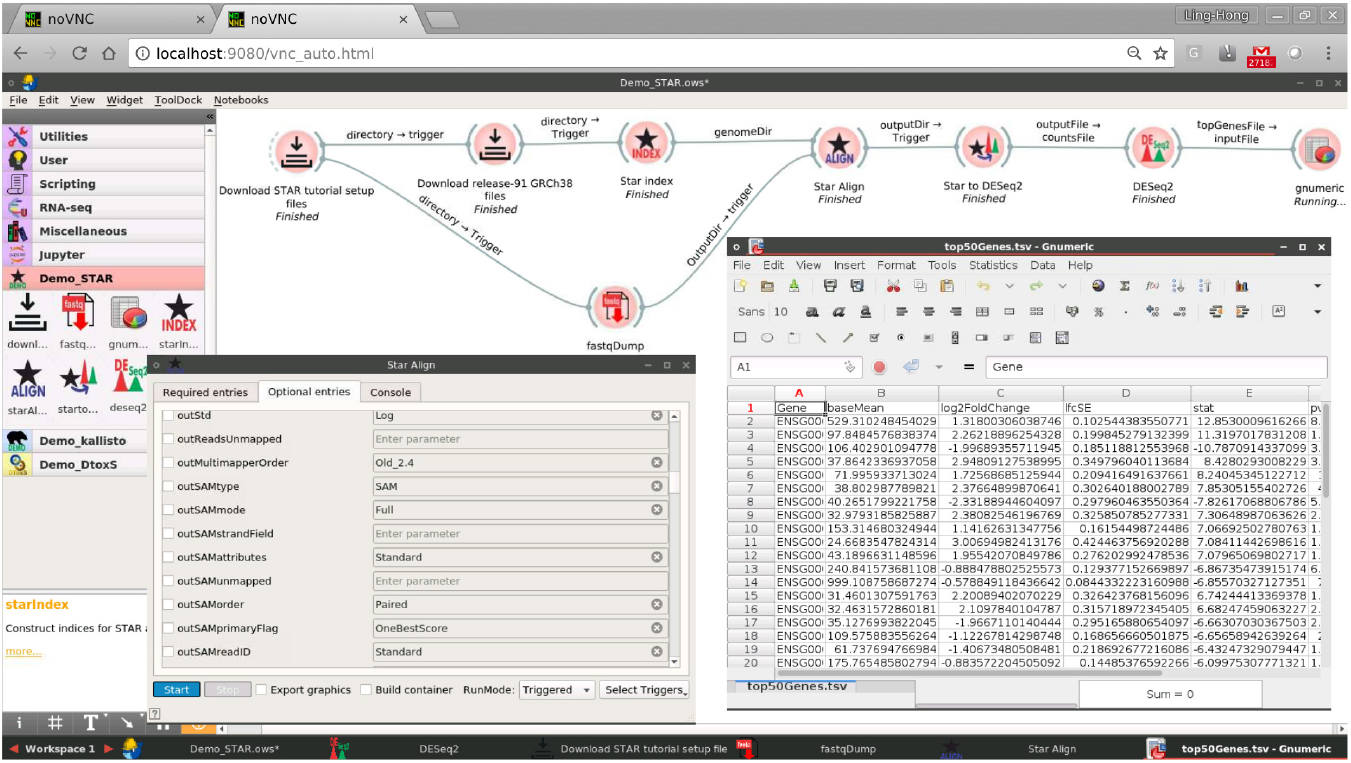
STAR/DESeq2 RNA-seq workflow on remote servers. The Bwb workflow consists of 9 widgets and implements a RNA-seq differential gene expression pipeline. This is very similar in structure to the Kallisto-sleuth pipeline. However, the STAR aligner requires more RAM (32 GB) than available on our laptop and the screenshot is from our browser connecting to the Bwb application running remotely on a local firewalled server. The connection is established using SSH tunneling. However, the gnumeric application pops up a window (lower right window) to view the final results and works identically when run remotely. The scrollable lower left window pops up upon clicking the STAR align widget. It shows some of the many parameters that the STAR aligner uses, that are carried along with the Bwb workflow and can be easily changed and customized.

#### Example 4: Using Jupyter notebooks and customizing a workflow

In this example, we illustrate the integration of Jupyter notebooks into Bwb workflows by using Jupyter notebook to analyze and visualize the data produced by kallisto. The notebook is a simplified version of the walkthrough from the Pachter lab [30, 31] and uses sleuth. One widget runs the nbconvert function to execute the Jupyter script. It then connects a second widget which opens the notebook automatically so that users can edit, interactively run the code and visualize results using the native Jupyter GUI. This demo is illustrated in Figure 6 where the notebook displays a box-plot of the expression of the top differentially expressed gene in the control and treatment samples. Using Bwb to incorporate Jupyter notebooks has many advantages. First, a Jupyter notebook can contain dynamic code for a single programming language, whereas a Bwb workflow can be constructed with many modules consisting of notebooks using different languages. Second, the use of containers ensures that the correct version of R is used with sleuth (3.4.4 and not 3.5.1.) and that the dependencies are pre-installed saving long installation times when running notebooks. Using Jupyter notebooks is an example method to customize Bwb workflows by allowing users to interactively modify the code and visualize results. More generally, widgets can be created and dropped into any workflow. We have included two videos as Additional File 2 and Additional File 3 involving this workflow. The video in Additional File 2 illustrates the steps involved in loading and executing this workflow. The video in Additional File 3 illustrates how to create a new widget that adds a custom Python script to trim the fastq files before analyses, and how to incorporate this new widget into the workflow.

**Figure 6.**
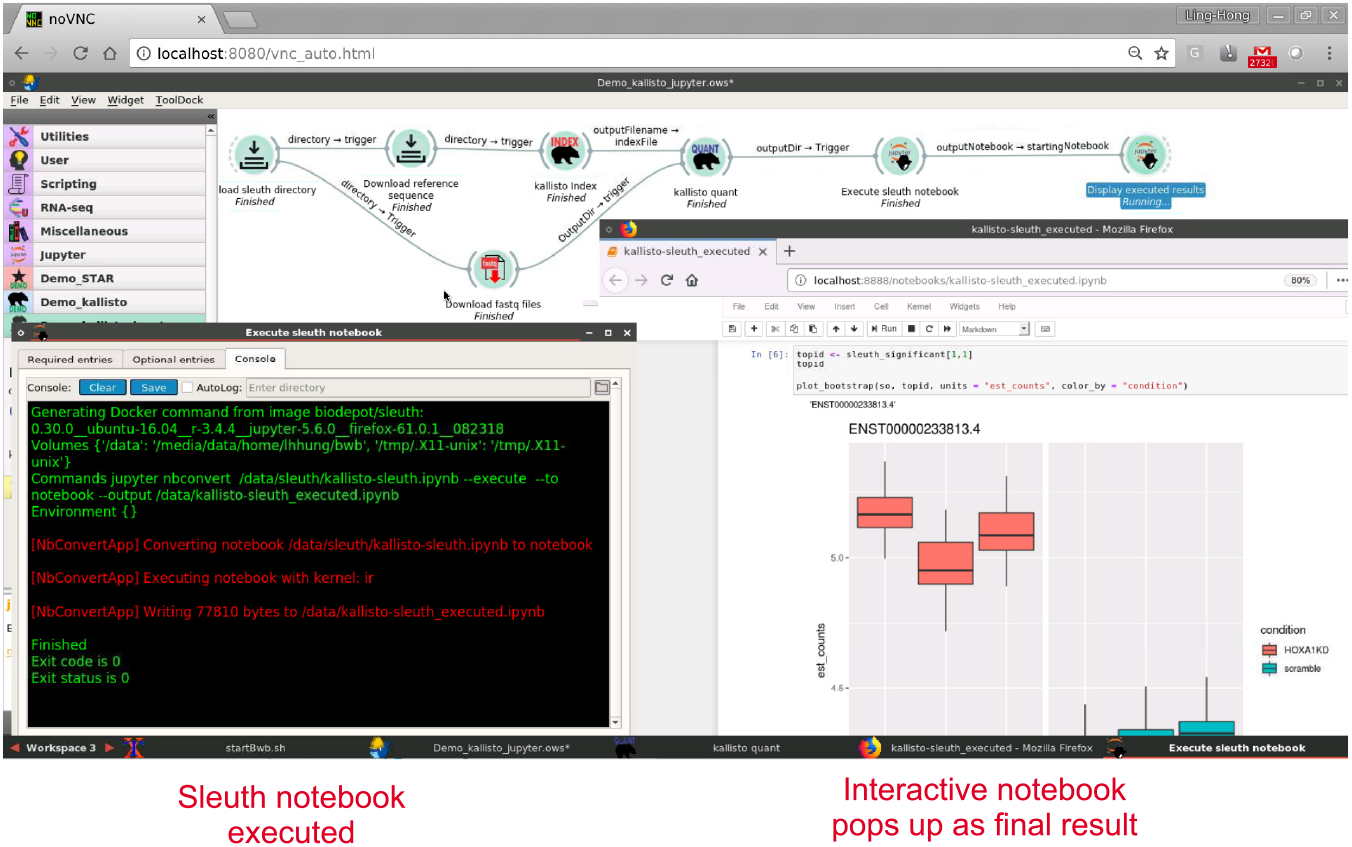
Kallisto with Jupyter-sleuth notebook: This workflow illustrates how simple it is to pop in different modules. In this case, a Jupyter notebook with a sleuth recipe to analyze the the kallisto data. The notebook itself is downloaded by the downloadURL utility at the beginning. The final two widgets execute and open the executed notebook. The left panel shows the console of the notebook being executed by calling the nbconvert function of Jupyter which creates a new notebook with the code cells executed. When finished it passes the name of the new notebook to the second instance of the Jupyter-sleuth widget which is configured to pop up the notebook. The notebook is accessed through a Firefox instance in the Jupyter container. The user can use the Firefox browser to fully interact with the container, including re-running the cells, adding cells and saving the notebook and results to their host computer

## Discussion

The examples that we have presented highlight example typical use cases for Bwb. The first use case is an important one – disseminating a protocol for an experiment so that researchers can easily replicate and validate their results. Traditional methods such as written SOP’s are not always effective no matter how detailed. Our example demonstrates how one can create a completely executable flowchart that runs the techniques and exposes the parameters and data. No installation of additional software is required and the entire SOP can be replicated and tested using a GUI.

The second and third use case demonstrates how Bwb can be used with standard workflows that are commonly used in bioinformatics. These well-established workflows have been documented using Jupyter notebooks. However, even with these protocols, modules requiring different computing environments and installation of these modules can be challenging. Bwb captures all the steps in the workflow, ensuring an independent computing environment through the use of Docker containers enhancing reproducibility. Each step is modifiable, with the parameters outside of the container, which promotes customization of the workflow.

The fourth use case demonstrates how Bwb’s support exporting GUIs can be utilized to add Jupyter notebooks to a workflow. In this manner, interactive customization of code and visualization of results can be added to any Bwb workflow. We showcase enhanced encapsulation, flexibility and reproducibility of Jupyter notebooks when integrated in Bwb workflows.

Bwb has some limitations. First, iteration through a set of parameters (e.g. a set of fastq files) and multi-threaded execution are currently possible using wrapper scripts inside the container. We are actively developing a more portable scheduler system to address these needs. Second, GUI support for the bash scripts depends on having Bwb open, or having either native X11 or X11 emulation through GUIDock. Furthermore, the bash scripts currently use the directory structure of the host machine that launches Bwb. We are working on adding additional tools to facilitate mapping of filenames for use on different machines. We are also working on converting the Bwb workflows into other workflow descriptors such as the Workflow Description Language (WDL) [40] or Common Workflow Language (CWL) [41] to enhance compatibility with other workflow execution engines such as and Seven Bridges and the Broad Institute’s Cromwell [42]. Finally, we have done as much as possible to containerize the pipelines and Bwb itself to isolate it from the host platform. However, the interaction with Bwb is not completely independent of the host, as it depends upon the installation of Docker and the support of HTML5 by the web browser to render the graphics. HTML5 is still a relatively new standard and browser support is still variable, but this should improve in the future. The manual provided in Additional File 1, guides the user through the installation steps for different platforms.

## Conclusions

The BioDepot-workflow-builder (Bwb) project builds upon many of the ongoing efforts to enhance the reproducibility of computational research using Docker containers. The user-friendly and open-source graphical user interface makes containerized workflows accessible to biomedical researchers who are not programmers. We have also provided extensive tools to manage parameters, facilitate user data input and the customization containers. The key aim of Bwb is to allow researchers with varying levels of technical skill to deploy, reproducibly execute and test alternative algorithms with confidence. In this manner, workflows and analytical results are made accessible to a large and varied set of users. Using Bwb, the biomedical community can quickly vet, implement, and share new technical advances in data analyses.

## Methods

### Bwb windowing environment

The Bwb container launches a mini-webserver on host that is accessed using the browser. The server uses fluxbox [43], a compact windows manager to provide a graphical user interface similar to Windows or the MacOS. Fluxbox provides full graphical support using X11 to render the graphics internally on the server. Bwb uses the GUIdock-X11 [44] system to allow containerized applications (such as Jupyter, gnumeric) to export their graphics and GUI to the server’s internal screen. The noVNC [45] protocol is then used to transfer the internally rendered screen to the user’s browser which draws the graphics on the in browser window using HTML5.

Minimizing or closing the startup Bwb window reveals the background screen. Right clicking on the background brings up an application menu. For the basic Bwb container, there are 3 menu options, the Bwb app, a terminal to enter system commands, and the quit container option. Fluxbox provides 4 separate workspaces that are available which act as independent screens. Multiple instances of Bwb can be launched simultaneously. Windows can be resized, minimized, maximized as in other windowing systems. Cut and paste is supported between windows inside the container. Support will be added to allow cut and paste with the host system.

### Drag-and-drop user interface: Widgets, Tool Dock and canvas

When Bwb is started, the Bwb application window pops up. On the left hand side of the application window is a tool box (Tool Dock) with multiple tabs (drawers) which contain different collections of widgets. Clicking on the tab expands the toolbox drawer to reveal the contents. Drawers are organized by function. Bwb comes with a set of 24 ready-to-use widgets. These are all linked to containers available on our BioDepot repository on Docker hub. Any workflows constructed with these widgets will automatically download the necessary containers the first time that they are run and require no installation. Users can also create their own drawers. A new drawer is created whenever a workflow is loaded. Also widgets can be added and removed and drawers removed using the ToolDock Editor available from the menu bar. To interact with a widget and include it in a workflow, the widget is dragged onto the canvas. Multiple copies of the same widget definition can exist in a workflow with different parameters. For example, the downloadURL widget is used twice in the kallisto demo workflows to download different files at different stages in the pipeline. Widgets on the canvas are then connected by dragging from the right side of the source widget to the left side of the sink widget. This will transfer the output of the source widget to the input of the sink widget during workflow execution.

The basic implementation of the Tool Dock and Canvas drag and drop UI is from OrangeML using the python PyQT5 Quicktime (QT) library. Orange workflows were connections of widgets that are given in the ToolDock, which once loaded, do not usually change. Bwb extends the Tool Dock to have drawers for each workflow to support custom widgets. Bwb provides the Tool Dock editor to allow for additional dynamic manipulations to the widgets in the Tool Dock. These additions are also implemented using PyQT5 and Python.

### Widget UI and definition windows

After being dragged to the Canvas, double clicking on the widget brings up a tabbed widget UI window, with tabs for entry of required and optional parameter values. A third tab reveals the console which displays intermediate results from execution. At the bottom of the window, a series of buttons and drop-down menus are available to control the execution of the widget.

Right clicking on the widget and choosing edit, brings up the tabbed widget definition window. Tabs are available for entry of values that define the UI window, choose port and volume mappings, inputs, outputs, the command to be executed, and the container to be used. In addition the user can access tools to build and manage containers using the Docker tab.

Bwb widgets are loosely based on widgets found in Orange. The functionality of Orange widgets reside in a Python script associated with the widget. With Bwb, the functionality resides in the commands, parameters and Docker container and not in the Python script. Accordingly Bwb stores these parameters in JSON files. A small Python script is still required in order to maintain compatibility with Orange routines such as the signal manager and widget manager that expect individual Python scripts. This stub Python script is auto-generated by Bwb’s widget builder module. The implementation of the forms in the UI and definition window is again accomplished using Python and PyQT5.

### Workflow storage and execution

The workflow is stored in a single directory. This directory contains widgets specific to the workflow, the icon, and a Python script used to load the workflow into Bwb. An XML file saves the connections between the widgets and all the parameter values or settings This is different information than the JSON files for each of the widgets which store information defining which parameters are queried and what and how the widgets execute based on these parameters.

Bwb takes the values from the widgets forms and generates a Docker command (or set of commands when there are multiple commands) for each widget. When the widget begins execution, a QT QProcess is launched. The QProcess runs the Docker command, prints the output to the console and checks for any signal sent by the user (through the ‘stop’ button) to abort the process. If the Docker command requires a container that is not present, Docker will automatically search through the public repositories and download the container if it is available (all containers used by widgets are available from the BioDepot repository). The QProcess then sends a signal when the Docker command has finished and a code to indicate whether the command successfully executed. If the Docker command was successful, the widget the sends out signals to downstream connected widgets. Upon receiving an input signal, the downstream widget checks that all necessary parameters are set and if execution is also triggered (or is automatic once parameters are set), its execution starts. In this manner, execution progresses down connected widgets in the workflow.

The Bwb signals are based on the Orange ML signals and are managed by the Orange signal manager. The XML connection files are also from OrangeML. The rest of the workflow storage and execution process is new to Bwb.

### Customizing workflows and containers

Customization of workflows can occur at different levels. The simplest is at the level of parameters. We have discussed how Bwb facilitates this through the use of forms and the separation of parameters from the Docker containers and through the use of interactive widgets such as Jupyter.

Customization can also be accomplished by replacing a module with another or adding an additional module such as a script. This would be accomplished by inserting a new widget into a workflow. Most modules will not have the exact same inputs and outputs, so this requires customization of widgets. We have already discussed how Bwb facilitates widget construction through its form based interface and provided a tutorial with a simple example.

Occasionally, the need will arise to customize Docker containers as well. Most bioinformatics scripts can be run within an off-the-shelf Bash, Python, Perl, R or Java containers that we have provided with Bwb. However, it is not uncommon to need to install libraries, and while these can be placed inside the script, it means that time consuming library installation steps using utilities like biocLite or install.packages must be executed each time the pipeline is run. To avoid this, we have provided our BiocImageBuilder tool [26] which allows users to specify packages required from Bioconductor and CRAN. BiocImageBuilder will build the container or provide a Dockerfile for the user to modify. In the future, we intend to expand the tool to include other common installation methodologies based on Conda and pip. Bwb also supplies files used to build its containers with its widgets in the Dock-erfiles directory, to allow the user to easily customize the images if they wish to do so without having to start from scratch. Our tutorial and video demonstrates how these tools are used a new container and widget for an existing script.

### Bwb code organization summary

The key components from OrangeML used by Bwb are the signal manager, Tool Dock and Canvas routines The signal manager queues and manages signals between widgets in workflows. The Tool Dock and Canvas modules handle the Tool Dock and the drag and drop interface. The OrangeML code has been forked and stored in a directory in the BioDepot-workflow-builder repository. Any modified Orange routines are kept in a separate directory.

The major Bwb modules are the BwBase, WidgetBuilder, ImageBuilder, Docker-Client, ToolDockEdit classes and WorkflowTools and CreateWidget packages. The BwBase class handles the widget UI window. The WidgetBuilder class is responsible for the widget definition window. The ImageBuilder class runs the BiocImageBuilder tool for building Docker containers. The ToolDockEdit class adds tools to edit the Tool Dock. These are all subclasses of the original orange widget class, largely to ensure that they interact correctly with the other OrangeML routines. Finally there are two new packages workflowTools and createWidget. WorkflowTools handles the loading and saving of Bwb workflows and the createWidget package creates JSON files and python files for the widgets.

## 1 Availability of source code and requirements

- Project name: BioDepot-workflow-builder (Bwb)
- Project home page: https://github.com/BioDepot/BioDepot-workflow-builder
- Contents available for download: Docker Images, Dockerfiles, installation scripts, execution scripts and demo videos.
- Operating system(s): Linux, Mac OSX, Microsoft Windows, Azure, AWS, Google Cloud Platform.
- Programming language(s): Python, HTML, JavaScript
- Other requirements: Docker version 1.13.1 or greater
- License: MIT License

## 2 Availability of supporting data and materials

**Additional File 1**: Bwb User Manual.

**Additional File 2**: Video of running Bwb, loading and executing the kallisto-sleuth-Jupyter RNA sequencing workflow (case study 4). Publicly available at https://youtu.be/jtu-jCU2DU0.

**Additional File 3**: Video of creating a new widget in Bwb that adds a custom Python script to trim the input fastq files before performing alignment using kallisto. This video also shows how to incorporate this new widget into the kallisto-sleuth RNA sequencing workflow.

## 3 Declarations

### 3.1

#### List of abbreviations

AWS: = Amazon Web Services
BWA: = Burroughs-Wheeler Aligner
Bwb: = BioDepot-workflow-Builder
DToxS: = Drug Toxicity Signature Generation Center
RNA-seq: = RNA sequencing
SOP: = standard operating procedure
VNC: = virtual network computing

### 3.4 Author’s Contributions

L.H.H is the primary developer for the Bwb project, wrote the existing code for the widgets and containers, adapted the OrangeML code for the drag and drop interface and toolbox, and implemented the Bwb container. K.Y.Y. coordinated the manuscript preparation. L.H.H. and K.Y.Y. drafted the manuscript. L.H.H. designed the framework of Bwb and the case studies. J.H. and T.M. contributed code to early Docker-py implementations of Bwb. J.H. designed and implemented the image building tool used by Bwb. L.H.H, J.H., T.M., D.K. and A.I. contributed to testing and case studies of the project. L.H.H., D.K., J.H. and K.Y.Y. contributed to the writing of the user manual. J.H., D.K. and L.H.H. made the videos in additional data files. L.H.H. and W.L. provided technical guidance to all the students. Y.X., E.U.A., M.R.B. and E.A.S. developed the computational analysis pipeline at DToxS. All authors tested Bwb, read and approved the final manuscript. …

## 4 Acknowledgements

We would like to thank Jayant Keswani and students who took TCSS 592 at University of Washington Tacoma for contributing widgets and testing effort for earlier versions of the Bwb project. We would also like to acknowledge the Student High Performance Computing Club and the eScience Institute, both at the University of Washington, for providing technical assistance and computing resources to Jiaming Hu….

---

Additional File 1: Bwb Manual https://github.com/BioDepot/BioDepot-workflow-builder

Additional File 2: Introductory tutorial on using Bwb https://www.youtube.com/watch?v=jtu-jCU2DU0

Additional File 3: Tutorial on adding a Python script to a Bwb workflow https://youtu.be/r_03_UG1mBg

